# Effect of growing regions on morphological characteristics, protein nutrition, rumen degradation and molecular structures of various whole-plant silage corn cultivars

**DOI:** 10.1101/2023.02.17.529020

**Authors:** Xinyue Zhang, Nazir Ahmad Khan, Enyue Yao, Fanlin Kong, Ming Chen, Rifat Ullah Khan, Xin Liu, Yonggen Zhang, Hangshu Xin, with the Lorem Ipsum Consortium

## Abstract

Little information exists on the variation of morphological characteristics, nutritional value, ruminal degradability, and molecular structural makeup of diverse whole-plant silage corn (WPSC) cultivars among different growing regions. This study investigated the between-regions discrepancies in five widely used WPSC cultivars in China (FKBN, YQ889, YQ23, DK301 and ZD958) in terms of 1) morphological characteristics; 2) crude protein (CP) chemical profile; 3) Cornell Net Carbohydrate and Protein System (CNCPS) CP subfractions; 4) *in situ* CP degradation kinetics; and 5) CP molecular structures. Our results revealed that significant difference were observed on growing region and WPSC cultivar interaction for all estimated morphological characteristics (*P* < 0.001), CP chemical profile (*P* < 0.001), CNCPS subfractions (*P* < 0.001) and CP molecular structural features (*P* < 0.05). Except ear weight (*P* = 0.18), all measured morphological characteristics varied among different growing regions (*P* < 0.001). Besides, WPSC cultivars planted in different areas had remarkably different (*P* < 0.01) CP chemical profiles and CNCPS subfractions. All spectral parameters of protein primary structure of WPSC differed (*P* < 0.05) due to the various growing regions, except amide II area (*P* = 0.28). Finally, the area ratio of amide I to II was negatively correlated with the contents of soluble CP (*δ* = -0.66; *P* = 0.002), CP (*δ* = - 0.61; *P* = 0.006), non-protein N (*δ* = -0.56; *P* = 0.004) and acid detergent insoluble CP (*δ* = - 0.43; *P* = 0.008), in conjunction with positively correlated with moderately degradable CP (PB_1_; *δ* = 0.58; *P* = 0.01). In conclusion, the current study suggested that even for the same WPSC cultivar, the morphological characteristics, protein nutritional values and rumen degradability significantly varied among different grown regions due to distinguished molecular structures.

**Author summary:** As the major roughage source, whole plant silage corn plays an essential role in ruminant feed industry. The quality and quantity of it largely influenced by environmental and climate conditions except genetic factors. However, there was limited information to systematically analyze whole plant silage corn from morphological characteristics, nutritional components, rumen degradation to its inherent molecular structures. Thus, this study was conducted to investigate the discrepancies of various silage-corn cultivars grown in different regions from internal structure to phenotype based novel technology - fourier transform infrared spectroscopy.

## Introduction

Globally, whole plant silage corn (WPSC) is a major ingredient in dairy total mixed ration (TMR) under most dietary regimes, owing to its high and stable biomass yield under a wide range of environment and soil conditions, in conjunction with its high contents of total digestible nutrients and metabolizable energy [1-3]. Moreover, the inclusion of corn silage in TMR increased dry matter (DM) and metabolizable energy intakes as well as milk yield of dairy cows [4]. Because of these great qualities, WPSC contributed up to 42% of TMR for high-producing dairy cows [5]. Thus, optimizing the biomass yield and nutritional qualities of WPSC is a long-term goal for dairy industry to meet the nutritional requirements of high production dairy cows on sustainable basis.

The exception of genotype and harvest maturity, the yield and nutritional quality of WPSC are highly influenced by environmental conditions [6-8]. For instance, high growing temperature reduced the digestibility of corn silage, which is a result of increased lignin content in stovers and decreased starch content in cobs [9-10]. A close association was observed between soil moisture and plants’ nutrients distribution in eight cultivars of napiergrass [10]. Moreover, Temime et al. [9] also reported that besides genetic factors, the proportion of volatile constituents in Chétouiolive oils was strongly influenced by growing regions. Similarly, the glycosidase inhibitory rate in mulberry leaves ranged from 142 to 845 g/kg between different cultivate regions were observed [11].

Although lower content of crude protein on average (CP; 74 g/kg DM) was observed in corn silage, it significantly contributed to the overall supply of metabolizable protein for dairy cows because of high consumption [3]. Moreover, rumen protein degradability and overall supply of metabolizable protein to dairy cows were strongly influenced by variations of CP chemical profiles and protein inherent molecular structures in feedstuffs [4, 12]. Recent studies suggested that the molecular structure of roughages could be revealed by advanced vibrational molecular spectroscopy techniques, for example, attenuated total reflectance - Fourier transform infrared spectroscopy (ATR-FTIR) [13-15]. Thus, it is possible to associate the protein chemical profiles and in situ degradation parameters with the specific inherent structures [16-19].

To our knowledge, no systematic study was conducted on morphological characteristics, CP rumen degradabilities and molecular structures of various WPSC cultivars planted in different growing regions. Therefore, in the present study, five widely utilized WPSC cultivars planted in five growing regions (Beijing, Urumchi, Cangzhou, Tianjin and Liaoyuan) in China were evaluated the differences on 1) morphological traits; 2) CP chemical profile; 3) Cornell Net Carbohydrate and Protein System (CNCPS) CP subfractions; 4) in situ CP degradation kinetics; and 5) protein molecular structures. The relationships between protein molecular structural parameters and CP chemical profile, CNCPS subfractions and ruminal degradation characteristics were also investigated. We hypothesized that the morphological characteristics, CP chemical profile, and ruminal degradability may differ in WPSC cultivated in different growing regions, and those discrepancies in protein nutritional value and rumen degradability were correlated with their inherent protein molecular structures.

## Materials and methods

### Corn cultivars, growing regions, and crop production

Five WPCS cultivars widely utilized for production in China were selected for this study, including Fengkenbainuo (FKBN), Yaqing 889 (YQ889), Yuqing 23 (YQ23), Dongke 301 (DK301) and Zhengdan 958 (ZD958). All samples were grown in five different regions, which were Beijing (39°9’ N, 116°3’ E), Urumchi (43°8’ N, 87°7’ E), Cangzhou (38°3’ N, 116°8’ E), Tianjin (39°1’ N, 117°2’ E) and Liaoyuan (42°7’ N, 125°5’ E). In each region, these five cultivars were sown in three plots (5 m × 5 m) with seeding rate of 1400 kernels per plot, in conjunction with plant-to-plant distance of 35 cm and row to row distance of 60 cm. All plots in five growing regions were treated under the same practical management and agronomic practices during cultivation. Data on geographical and climate condition of five experimental regions were summarized in Table 1.

**Table 1.** Longitude and latitude, average annual temperature, and average annual precipitation of different growing regions of whole-plant silage corn cultivars.

| Region | Longitude | Latitude | Temperature (°C) | Precipitation (mm) |
| --- | --- | --- | --- | --- |
| *Beijing* | E 116°3' | N 39°9' | July: 23~31 | July: 97.5 |
|  |  |  | August: 21~30 | August: 233.9 |
|  |  |  | September: 17~27 | September: 2.8 |
|  |  |  | October: 7~15 | October: 73.3 |
| *Urumchi* | E 87°7' | N 43°8' | July: 21~31 | July: 3.2 |
|  |  |  | August: 17~26 | August: 14.5 |
|  |  |  | September: 11~22 | September: 3.7 |
|  |  |  | October: 2~11 | October: 8.8 |
| *Cangzhou* | E 116°8' | N 38°3' | July: 24~32 | July: 161.0 |
|  |  |  | August: 22~30 | August: 124.7 |
|  |  |  | September: 18~29 | September: 4.7 |
|  |  |  | October: 8~16 | October: 138.6 |
| *Liaoyuan* | E 125°5' | N 42°7' | July: 19~30 | July: 231.8 |
|  |  |  | August: 17~27 | August: 259.0 |
|  |  |  | September: 9~23 | September: 16.7 |
|  |  |  | October: 0~12 | October: 1.6 |
| *Tianjin* | E 117°2' | N 39°1' | July: 25~32 | July: 159.6 |
|  |  |  | August: 23~31 | August: 160.4 |
|  |  |  | September: 19~29 | September: 1.2 |
|  |  |  | October: 9~17 | October: 101.9 |

### Crop sampling and morphological measurements

Twenty plants (10 cm above the ground) from each plot were randomly harvested at kernel maturity stage of half milk line when targeted DM content was around 330 g/kg. The exterior 1 m area of each plot was excluded from sampling to ensure uniformity in the sampled plants. The physical parameters of WPCS including whole-plant height, whole-plant weight, stem diameter, ear height, ear weight, ear length, ear diameter, ear rows and kernel numbers in each row were measured and recorded at harvest. Then, the whole-plant samples were chopped to small particle size (1-2 cm), dried (at 60 °C for 24 h) and grounded through a 1.0 mm screen (FZ102, Taisite Instrument Co., Ltd, Tianjin, China) for analysis of chemical components and CNCPS compositions, or through a 2 mm screen for *in situ* incubation measurements, or through a 0.25 mm screen for molecular spectral analysis of protein primary and secondary structures.

### Protein chemical analysis and subfractions composition

The WPSC samples were analyzed for CP content (official method 984.13, using a Kjeltec 2400 autoanalyzer; Foss Analytical A/S, Hillerϕd, Denmark) according to AOAC [20]. The contents of soluble CP (SCP), neutral detergent insoluble CP (NDICP) and acid detergent insoluble CP (ADICP) were determined according to Licitra et al [21].

For estimating the difference of CP degradation in the rumen, the CP subfractions composition were determined according to the updated version (v 6.5) of Cornell Net Carbohydrate Protein System [CNCPS, 22-23]. A total of five subfractions quantified in this study were: PA_1_ (non-protein N; ruminal degradation rate (k_d_) of 2/h), PA_2_ (soluble non-ammonia protein; k_d_ of 0.10-0.40/h), PB_1_ (moderately degradable CP; k_d_ of 0.03-0.20/h), PB_2_ (slowly degradable CP; k_d_ of 0.01-0.18/h) and PC (unavailable CP).

These CP subfractions were estimated from the chemical profiles according to the following mathematical models:

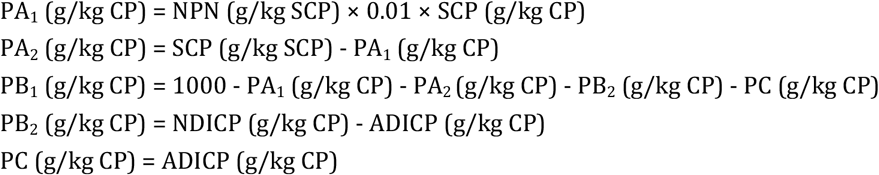

### *In situ* rumen degradation

Three healthy and non-lactating Holstein Frisian dairy cows (550-600 kg body weight) fixed with permanent rumen fistula from the experimental base of Northeast Agricultural University were used for *in situ* incubation research. These animals were fed TMR with forage to concentrate ratio of 60:40, twice daily at 07:00 and 21:00 h. The TMR was composed of 428 g/kg Chinese wildrye, 158 g/kg corn silage, 132 g/kg corn, 74 g/kg corn gluten feed, 54 g/kg dried distiller grains, 49 g/kg corn germ meal, 37 g/kg wheat bran, 32 g/kg soybean meal, 21 g/kg cottonseed meal, 10 g/kg molasses beet, in conjunction with 5 g/kg minerals and vitamins.

Subsamples (ca. 7 g) were randomly incubated in sealed nylon bags (10 × 20 cm, pore size 40 μm) in rumen of three fistulated cows for 0, 4, 8, 12, 24, 36, 48 and 72 h, using the “gradual in/all out” schedule. Three replicate bags per sample from individual cow were used for every incubation time point. After incubation, all nylon bags were taken out of the rumen, washed with cold running tap water for six times, and then dried to constant weight in forced air over at 65 °C. The dried residues from replicate bags of each sample were pooled according to the incubation time, then grounded and stored in plastic sealed bags for further analysis. Total three replicates were gained from three dairy cows. The parameters related to rumen degradation characteristics of CP were calculated according to Ørskov and McDonald [24].

### Spectral data collection and analysis

Data on protein primary and secondary structures were collected at the Chemical Molecular Structure Analysis Laboratory located in Northeast Agriculture University (Harbin, China). Fourier transform infrared spectroscopy (FT/IR; Bruker ALPHA-T, Germany) was utilized for the molecular spectral analysis. The spectral data were obtained at the mid-infrared fingerprint region from ca. 4000 to 400 cm^-1^, with 128 coadded scans at a spectral resolution of 4 cm^-1^. Each sample was scanned five times. Data were collected at peak height and area of the spectral bands related to protein primary structural functional groups. Only the spectral intensities of amide I band (baseline: ca. 1712 - 1565 cm^-1^; peak height: ca. 1639 cm^-1^) and amide II band (baseline: ca. 1565 - 1489 cm^-1^; peak height: ca. 1545 cm^-1^) were analyzed. The fourier self-deconvolution (FSD) method and the second derivative function in OMNIC 8.2 software (Spectra Tech., Madison, WI, USA) were used to characterize the protein secondary structure in amide I region including protein *α*-helix (peak height: ca. 1656 cm^-1^) and protein *β*-sheet (peak height: ca. 1634 cm^-1^). The protein molecular structure spectral data quantified in this study were the peak height and area of protein amide I, amide II and their ratio, in conjunction with peak height of *α*-helix, *β*-sheet and their ratio.

To visualize the structural differences in protein structures, multivariate analysis, agglomerative hierarchical cluster analysis (CLA) and principal component analysis (PCA) were applied on the overall spectroscopic data within the protein amide I and amide II region (ca. 1712 - 1489 cm^−1^), using Statistica 8.0 software (StatSoft Inc., Tulsa, OK, USA) [25].

### Statistical analysis

The PROC MIXED procedure of SAS 9.4 was used to analyze data on the plant morphological characteristics, CP chemical profiles, CNCPS subfractions, *in situ* CP degradation kinetics and protein molecular structural parameters, the model used was: Y_ijkl_ = µ + C_i_ + R_j_ + C_i_ × R_j_ + D_k_ +e_ijkl_, where Y_ijkl_ was an observation on the dependent variable ijkl, µ was the population mean of the variable, C_i_ was a fixed effect of corn cultivar (i = 5; FKBN, YQ889, YQ23, DK301 and ZD958), R_j_ was a fixed effect of growing region (j = 5; Beijing, Urumchi, Cangzhou, Tianjin and Liaoyuan), C_i_ × R_j_ was a fixed effect of interaction between factor C at level i and factor R at level j, D_k_ was the random effect of dairy cow, and eijkl was the model error.

The correlation between protein spectral parameters and protein chemical profile, CNCPS CP subfractions was well as *in situ* CP degradation data were analyzed by the Hmisc and Corrplot packages of R (version 4.0.2; R Foundation for Statistical Computing, Vienna, Austria). For all statistical analysis, significance was declared at *P* < 0.05, and tendency was declared at 0.05 < *P* < 0.10.

## Results and discussion

### Effects of growing regions and cultivars on morphologicalcharacteristics of whole plant silage corn

The significant interactions between cultivar and growing region on morphologgical characteristics were showed in Table 2 (*P* < 0.001). Except ear weight (*P* = 0.18), all measured morphological characteristics varied (*P* < 0.001) among the growing regions. Plant height and weight, in conjunction with ear height and length were highest (*P* < 0.05) in WPSC cultivars grown in Urumchi, while whole plant height and weight, as well as ear length were lowest (*P* < 0.05) in Tianjin. Cultivars grown in Beijing represented greatest (*P* < 0.05) stem diameter, ear diameter and rows of kernel, whereas Urumchi showed the opposite condition (*P* < 0.05). Mean number of kernels in a row was highest (*P* < 0.05) in Tianjin and lowest (*P* < 0.05) in Beijing. Previous studies have reported that precipitation was one of the most influential abiotic factors for plant productivity [26], and drought stress generally contributed to delay in plant growth and development by decreasing cell elongation and reducing photosynthesis [27]. Moreover, soil moisture and growing temperature were highly related to DM yields because they affected canopy and anatomical development of maize crop [10, 28]. Wang et al. [29] reported that the WPSC cultivar of Yunuo_7 had better leaf appearance rate at high growing temperature (37.5 °C, southwest of China) than other maize cultivars adopted to northeast and north plain of China. Furthermore, there was an effect of solar radiation on growth components which were proportional to gross photosynthesis [30]. Results of Tollenaar et al. [31] showed that more than a quarter of DM yield of maize crop was attributable to solar brightening during 1984 to 2013 in the US Corn Belt. In addition, limited solar radiation during flowering reduced kernel number and kernel development of maize [32]. Above factors may partially explain the reasons for differences in morphological characteristics of WPSC cultivars from different growing regions observed in our study.

**Table 2.** Morphological measurements of silage-corn cultivars grown in different regions. %.

| Cultivars | Regions | Whole plant |  | Stem | Ear |  |  |  |  | Row |
| --- | --- | --- | --- | --- | --- | --- | --- | --- | --- | --- |
|  |  | Height | Weight | Diameter | Height | Weight | Length | Diameter | Rows | kernels |
|  |  | (cm) | (kg) | (cm) | (cm) | (kg) | (cm) | (cm) |  |  |
| FKBN | Beijing | 340.0 | 1.35 | 2.31 | 156.2 | 0.53 | 31.2 | 8.26 | 17.2 | 28.6 |
|  | Urumchi | 320.1 | 2.26 | 1.66 | 168.7 | 0.84 | 29.7 | 5.72 | 13.8 | 40.8 |
|  | Cangzhou | 317.8 | 0.89 | 1.80 | 151.3 | 0.33 | 27.5 | 5.59 | 13.2 | 28.8 |
|  | Tianjin | 297.7 | 0.86 | 1.56 | 151.3 | 0.39 | 24.3 | 5.37 | 14.8 | 39.2 |
|  | Liaoyuan | 293.2 | 1.01 | 2.00 | 149.8 | 0.43 | 26.6 | 6.28 | 16.0 | 42.2 |
| YQ889 | Beijing | 316.0 | 0.96 | 2.09 | 152.5 | 0.36 | 24.4 | 5.92 | 16.6 | 30.1 |
|  | Urumchi | 363.6 | 1.17 | 1.84 | 190.8 | 0.35 | 28.0 | 5.89 | 15.2 | 30.6 |
|  | Cangzhou | 328.0 | 1.31 | 1.93 | 155.1 | 0.50 | 27.1 | 6.16 | 15.4 | 30.7 |
|  | Tianjin | 254.2 | 1.99 | 1.75 | 122.0 | 0.72 | 23.5 | 5.96 | 15.6 | 33.6 |
|  | Liaoyuan | 269.5 | 0.77 | 1.90 | 127.3 | 0.33 | 24.9 | 5.84 | 15.6 | 31.8 |
| YQ23 | Beijing | 280.4 | 1.78 | 2.20 | 124.7 | 0.83 | 24.8 | 6.33 | 16.4 | 34.9 |
|  | Urumchi | 345.5 | 0.96 | 1.59 | 186.3 | 0.31 | 29.3 | 5.64 | 13.8 | 24.2 |
|  | Cangzhou | 330.4 | 1.15 | 1.68 | 154.8 | 0.44 | 25.1 | 5.86 | 16.2 | 32.8 |
|  | Tianjin | 262.9 | 0.66 | 1.79 | 143.2 | 0.26 | 23.3 | 5.68 | 14.8 | 32.1 |
|  | Liaoyuan | 296.0 | 1.21 | 2.12 | 154.0 | 0.44 | 26.1 | 6.04 | 14.4 | 37.8 |
| DK301 | Beijing | 301.6 | 0.87 | 2.01 | 142.4 | 0.33 | 23.2 | 6.28 | 17.2 | 27.4 |
|  | Urumchi | 351.0 | 1.32 | 1.97 | 182.8 | 0.42 | 31.6 | 6.34 | 14.0 | 33.7 |
|  | Cangzhou | 282.9 | 1.93 | 1.71 | 131.8 | 0.80 | 23.4 | 6.24 | 14.6 | 32.2 |
|  | Tianjin | 257.6 | 0.88 | 1.98 | 124.0 | 0.41 | 24.7 | 6.42 | 16.0 | 39.8 |
|  | Liaoyuan | 298.6 | 1.02 | 1.69 | 158.1 | 0.33 | 27.3 | 5.72 | 14.8 | 34.8 |
| *ZD958* | *Beijing* | 271.8 | 0.80 | 1.87 | 137.6 | 0.33 | 24.0 | 6.15 | 15.8 | 26.7 |
|  | *Urumchi* | 383.3 | 1.19 | 1.62 | 208.9 | 0.34 | 28.1 | 4.85 | 13.8 | 36.0 |
|  | *Cangzhou* | 301.4 | 1.28 | 2.00 | 137.4 | 0.46 | 26.2 | 6.47 | 14.0 | 36.5 |
|  | *Tianjin* | 340.5 | 1.08 | 2.32 | 185.3 | 0.53 | 23.9 | 6.41 | 17.7 | 40.3 |
|  | *Liaoyuan* | 260.7 | 1.76 | 1.84 | 117.8 | 0.72 | 24.3 | 6.00 | 14.8 | 37.1 |
|  | *SEM* | 69.04 | 0.82 | 0.42 | 46.54 | 0.34 | 4.85 | 1.16 | 2.36 | 9.36 |
| *Regions* |  |  |  |  |  |  |  |  |  |  |
|  | *Beijing* | 301.96<sup>c</sup> | 1.15<sup>b</sup> | 2.10<sup>a</sup> | 142.68<sub>b</sub> | 0.48<sup>ab</sup> | 25.52<sup>b</sup> | 6.59<sup>a</sup> | 16.64<sup>a</sup> | 29.54<sup>c</sup> |
|  | *Urumchi* | 352.70<sup>a</sup> | 1.38<sup>a</sup> | 1.74<sup>c</sup> | 187.50<sup>a</sup> | 0.45<sup>b</sup> | 29.34<sup>a</sup> | 5.69<sup>c</sup> | 14.12<sup>d</sup> | 33.06<sup>b</sup> |
|  | *Cangzhou* | 312.10<sub>b</sub> | 1.31<sup>a</sup> | 1.82<sup>bc</sup> | 146.08<sub>b</sub> | 0.51<sup>a</sup> | 25.86<sup>b</sup> | 6.06<sup>b</sup> | 14.68<sup>cd</sup> | 32.20<sup>b</sup> |
|  | *Tianjin* | 282.58<sub>d</sub> | 1.09<sup>b</sup> | 1.88<sup>b</sup> | 145.16<sub>b</sub> | 0.46<sup>ab</sup> | 23.94<sup>c</sup> | 5.97<sup>b</sup> | 15.78<sup>b</sup> | 37.00<sup>a</sup> |
|  | *Liaoyuan* | 283.60<sub>d</sub> | 1.15<sup>b</sup> | 1.91<sup>bc</sup> | 141.40<sub>b</sub> | 0.45<sup>b</sup> | 25.84<sup>b</sup> | 5.98<sup>b</sup> | 15.12<sup>bc</sup> | 36.74<sup>a</sup> |
|  | *SEM* | 24.71 | 0.44 | 0.38 | 18.75 | 0.19 | 3.60 | 0.71 | 2.45 | 8.81 |
| *P-value* | *Regions* | <0.001 | <0.001 | <0.001 | <0.001 | 0.18 | <0.001 | <0.001 | <0.001 | <0.001 |
|  | *Cultivar* | <0.001 | 0.38 | 0.75 | 0.002 | 0.29 | <0.001 | 0.001 | 0.35 | 0.001 |
|  | *Region × Cultivar* | <0.001 | <0.001 | <0.001 | <0.001 | <0.001 | <0.001 | <0.001 | <0.001 | <0.001 |

Note: FKBN = Fengkenbainuo; YQ 889 = Yaqing 889; YQ 23 = Yuqing 23; DK 301 = Dongke 301; ZD958 = Zhengdan 958.

### Effects of growing regions and cultivars on protein chemical profiles and CNCPS of whole plant silage corn

The CNCPS was widely used for the evaluation of ruminant feedstuff [33-36]. In the present study, significant effect of cultivar and growing region interaction (*P* < 0.001) was observed for all measured CP chemical profiles and CNCPS subfractions (Table 3). WPSC cultivars grown in different areas exhibited remarkably different CP chemical profiles. For instance, the concentrations of ADICP (3.6 g/kg) and PC (49.3 g/kg) were lowest (*P* < 0.05) in WPSC cultivated in Urumchi. The results suggested that WPSC grown in Urumchi contained high proportion of protein available to animals, and implied that climate conditions (e.g. precipitation and growing temperature) could influence internal nutrient accumulations of WPSC. In agreement with our findings, previous study reported that the CP contents in maize and wheat were lower in rainy years [37]. High growth temperature markedly increased the lignin content of maize plants [11, 38], which can partly explain the lowest contents of ADICP (3.6 g/kg) and PC (49.3 g/kg) cultivated in Urumchi, where the lowest temperature was recorded among all five growing regions. As for the soluble protein contents, such as NPN and SCP, the WPSC planted in Beijing had higher (*P* < 0.05) content than other regions. Similarly, the CP content was greatest in Beijing (86.6 g/kg), followed by Tianjin (85.7 g/kg), Cangzhou (78.2 g/kg), Liaoyuan (78.0 g/kg) and Urumchi (73.5 g/kg). In terms of cultivar effect, protein content varied among WPSC cultivars. The highest CP content was recorded for DK301 (85.6 g/kg) and lowest for ZD958 (74.4 g/kg). In addition, the greatest variation range of CP among different regions was found for YQ23 cultivar (69.0 to 90.9 g/kg) and the lowest for DK301 (80.6 to 91.4 g/kg).

**Table 3.**
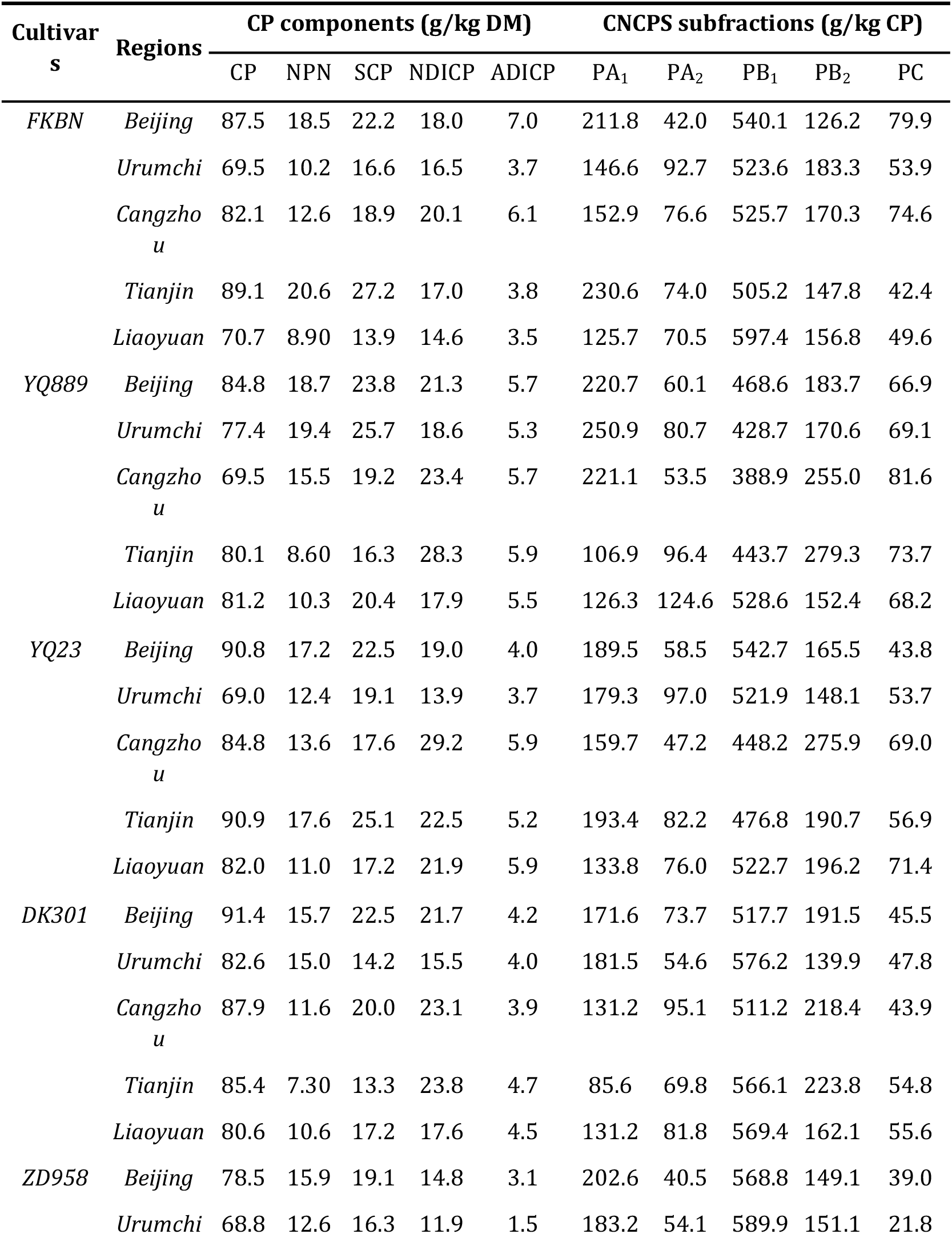
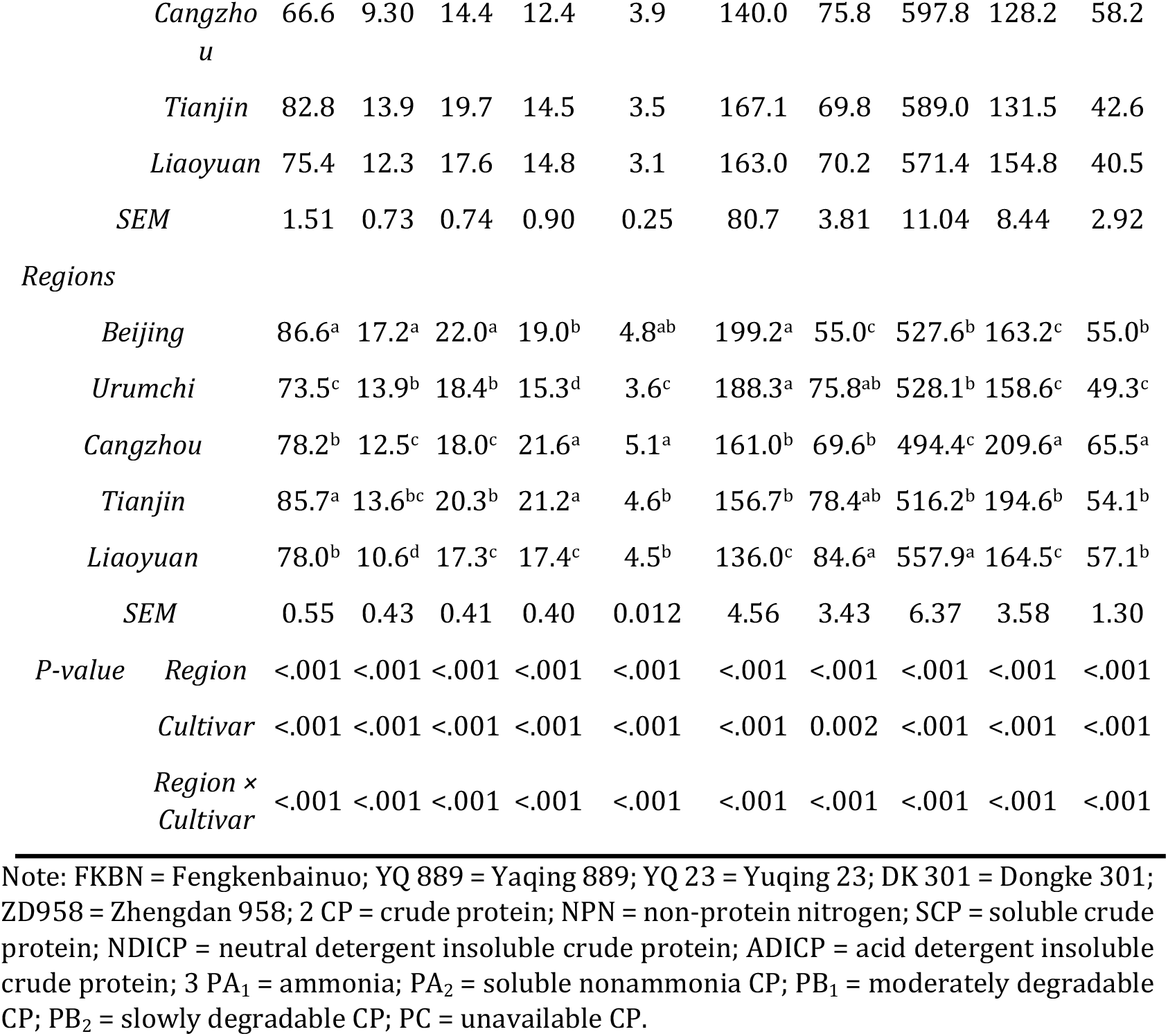
Crude protein (CP) chemical components and the CNCPS subfractions of whole plant silage corn cultivars grown in different regions. %.

### Effects of growing regions and cultivars on protein *in situ* rumen degradation of whole plant silage corn

A significant interaction of corn cultivar and growing region was observed in *in situ* soluble CP (S; *P* = 0.002) and ruminal degradable CP (RDP; *P* < 0.001). However, the interactive effect was not significant (*P* > 0.05) in the rate of degradation (K_d_), potentially ruminal degradable fraction CP (D), undegradable CP (U), and ruminal undegradable CP (RUP) (Table 4). Among *in situ* CP degradable characteristics, only the content of RUP varied (*P* < 0.001) among growing regions (Table 4). The highest (*P* < 0.05) content of RUP was observed in the WPSC grown in Beijing (41.7 g/kg DM), followed by Tianjin (39.3 g/kg DM), Liaoyuan (34.9 g/kg DM), Urumchi (34.7 g/kg DM) and Cangzhou (33.0 g/kg DM). In terms of cultivars, the highest (*P* < 0.05) content of RDP was observed in DK301 (47.8 g/kg DM) and lowest in ZD958 (40.2 g/kg DM). However, the highest and lowest variation of RDP content among regions was FKBN (33.7 g/kg DM in Urumchi to 53.2 g/kg DM in Tianjin) and YQ889 (38.3 g/kg DM in Cangzhou to 44.5 g/kg DM in Beijing), respectively. Cox et al. [39] reported that *in vitro* NDF digestibility of six corn hybrids on average varied from 710 to 827 g/kg between two growing regions, which indirectly explained our results. But limited information exists to make a comparison of *in situ* CP rumen degradation kinetics for forages cultivated in different regions. The variation of RDP in WPSC from different regions was highest in FKBN (39.3 to 53.2 g/kg DM), followed by YQ23 (38.3 to 46.5 g/kg DM), DK301 (43.5 to 47.9 g/kg DM), ZD958 (36.0 to 44.9 g/kg DM) and YQ889 (39.8-44.5 g/kg DM). The result was not fully consistent with the variation range observed in CP content.

**Table 4.**
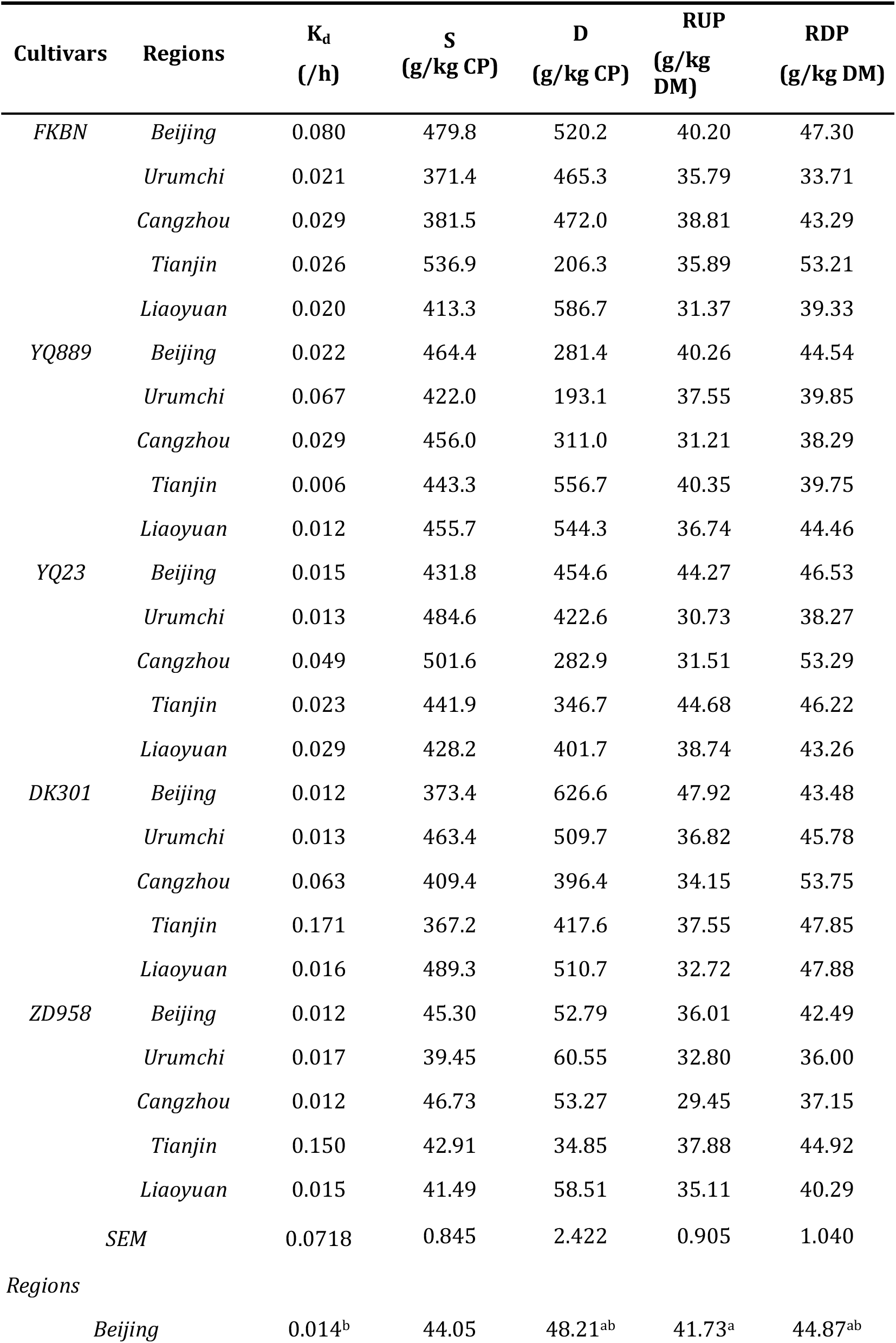
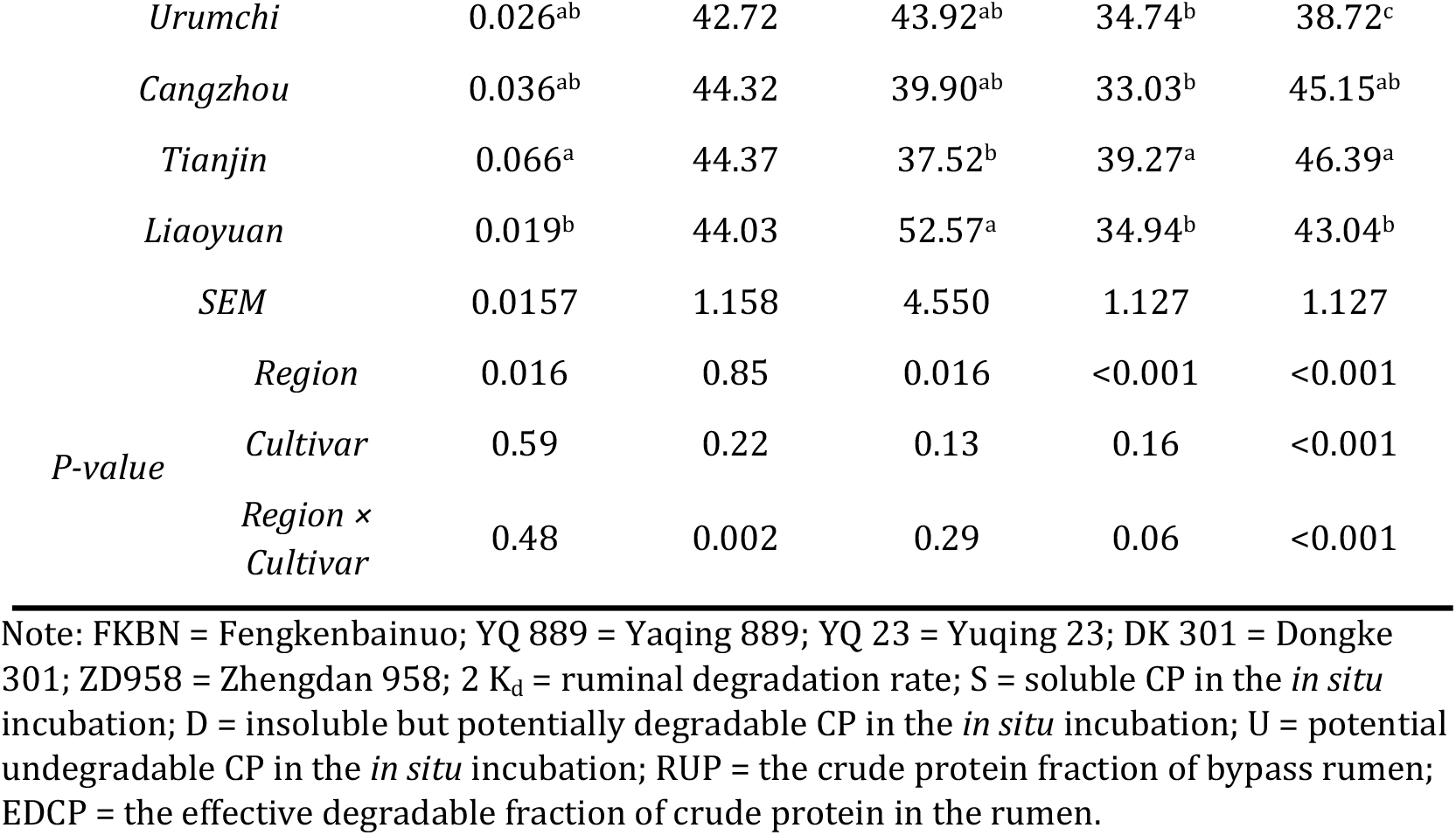
In situ crude protein (CP) degradation kinetics of whole plant silage corn cultivars grown in different regions. %.

### Effects of growing regions and cultivars on protein molecular structures of whole plant silage corn

Previous studies showed that feed protein solubility, ruminal degradability and post-ruminal digestibility not only depended on CP chemical compositions, but also strongly influenced by protein inherent molecular structures [40-42]. Therefore, differences in protein spectral characteristics of WPSC were quantified and visualized between growing regions and corn cultivars.

Data on secondary protein structures showed an interactive effect of growing region and cultivar for *α*-helix and *β*-sheet height (Table 5, *P* < 0.001). However, no significant interaction was observed on the *α*-helix to *β*-sheet height ratio (*P* = 0.26). The ration of *α*-helix to *β*-sheet height varied (*P* < 0.001) among growing regions, specifically, the highest ratio was recorded for Urumchi (1.88) and lowest for Beijing (1.02). In agreement with our findings, previous study reported that crops under water stress exhibited significantly different CP structural features and carbohydrates spectral regions compared to crops grown in dry land [6, 8]. During the growing and development period, the plants need to absorb and accumulate nutrients by the roots from soil, and water is the key transmission medium for transportation of all nutrients. Therefore, sufficient soil water could contribute to high efficiency of nutrients transportation and promotion of cell elongation [43], which might further cause a great development of molecular makeup and conformation of biopolymers in plants. The highest variation in protein amide area across the growing regions was observed in YQ889 (2.5 to 11.7 units) and lowest in DK301 (5.4 to 9.0 units).

**Table 5.**
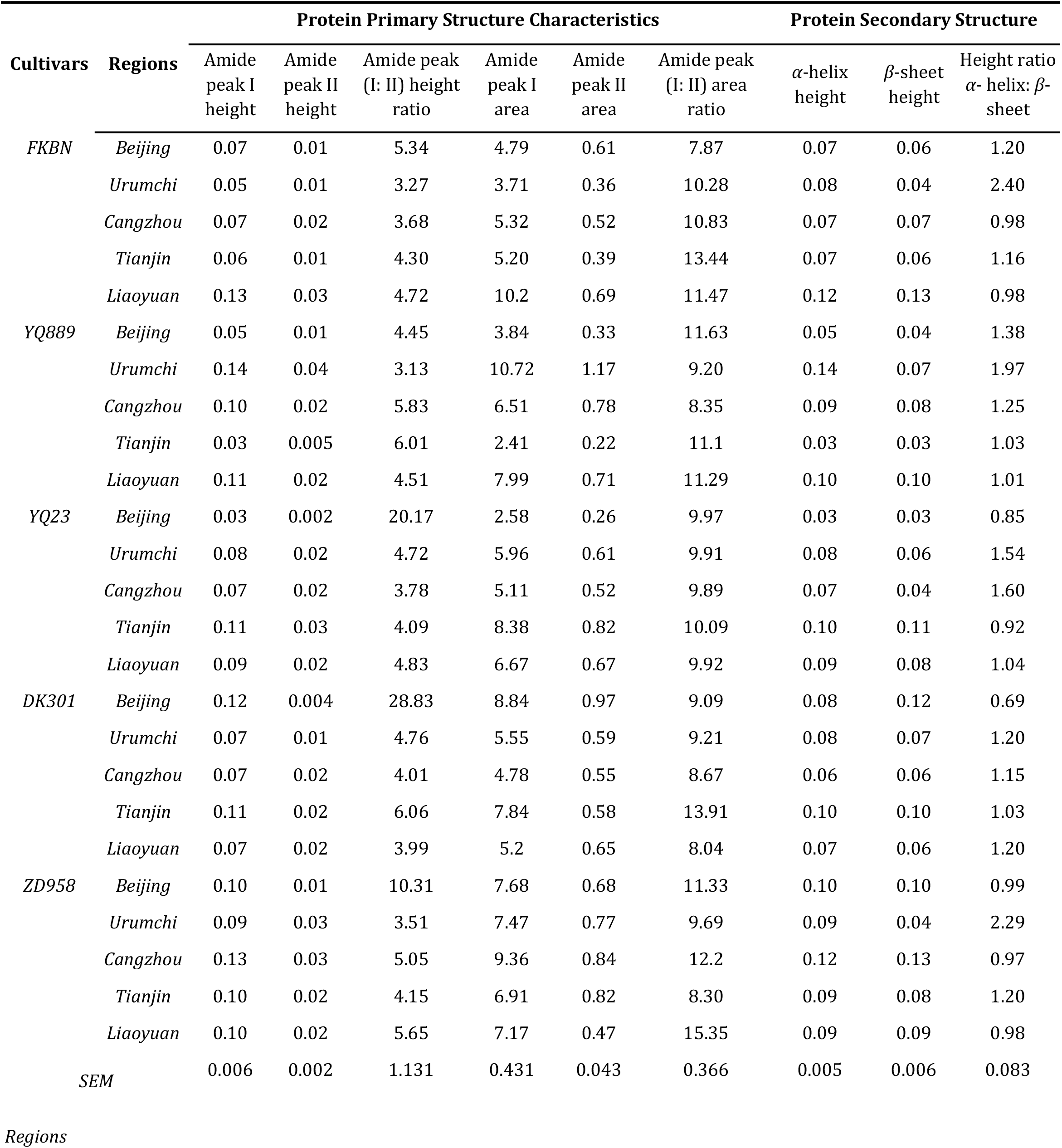
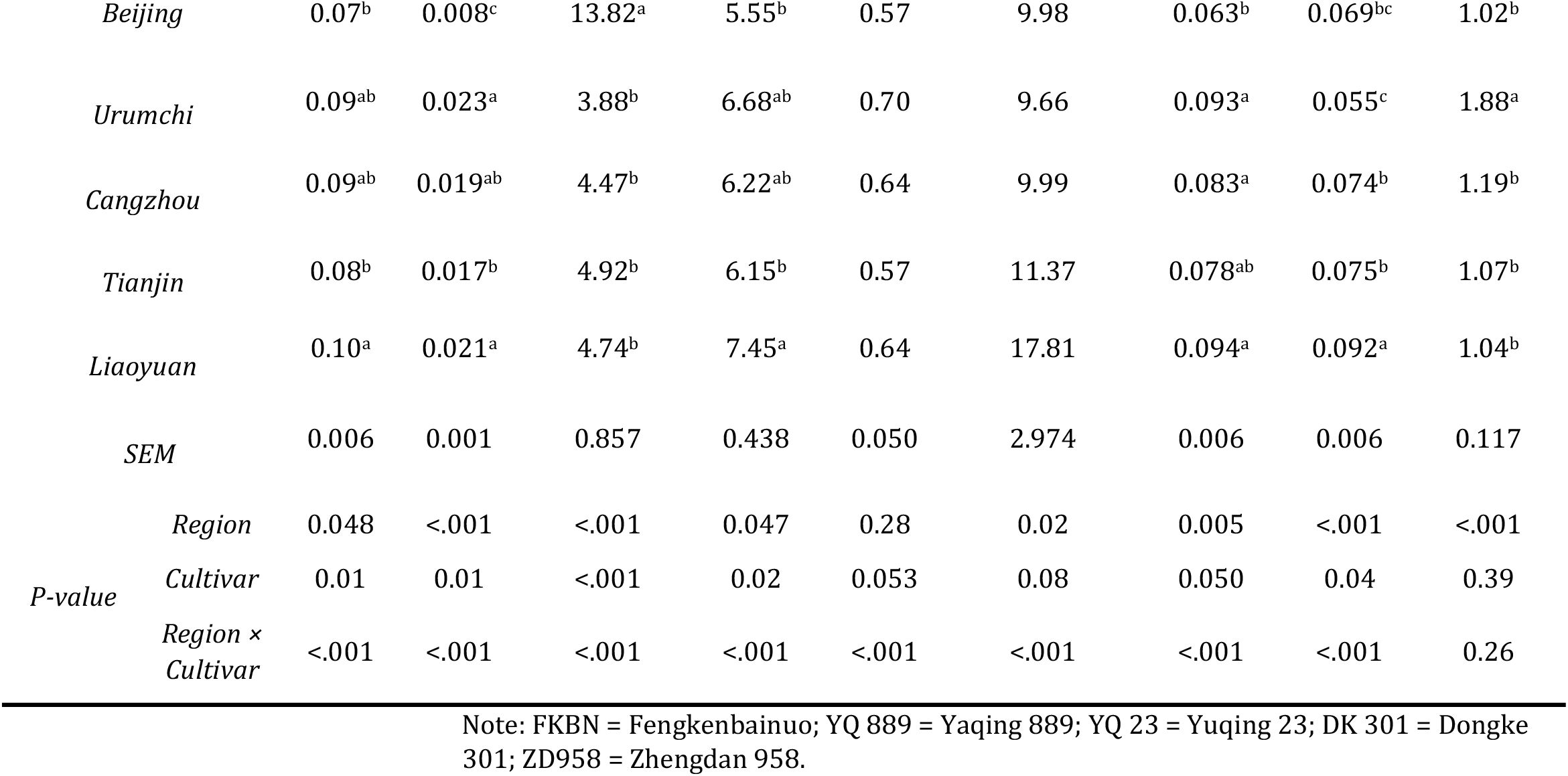
Protein molecular structures of silage corn cultivars grown in different regions. %.

| Cultivars | Regions | Protein Primary Structure Characteristics |  |  |  |  |  | Protein Secondary Structure |  |  |
| --- | --- | --- | --- | --- | --- | --- | --- | --- | --- | --- |
| | | Amide peak I height | Amide peak II height | Amide peak (I: II) height ratio | Amide peak I area | Amide peak II area | Amide peak (I: II) area ratio | α-helix height | β-sheet height | Height ratio α-helix: β-sheet |
| FKBN | Beijing | 0.07 | 0.01 | 5.34 | 4.79 | 0.61 | 7.87 | 0.07 | 0.06 | 1.20 |
|  | Urumchi | 0.05 | 0.01 | 3.27 | 3.71 | 0.36 | 10.28 | 0.08 | 0.04 | 2.40 |
|  | Cangzhou | 0.07 | 0.02 | 3.68 | 5.32 | 0.52 | 10.83 | 0.07 | 0.07 | 0.98 |
|  | Tianjin | 0.06 | 0.01 | 4.30 | 5.20 | 0.39 | 13.44 | 0.07 | 0.06 | 1.16 |
|  | Liaoyuan | 0.13 | 0.03 | 4.72 | 10.2 | 0.69 | 11.47 | 0.12 | 0.13 | 0.98 |
| YQ889 | Beijing | 0.05 | 0.01 | 4.45 | 3.84 | 0.33 | 11.63 | 0.05 | 0.04 | 1.38 |
|  | Urumchi | 0.14 | 0.04 | 3.13 | 10.72 | 1.17 | 9.20 | 0.14 | 0.07 | 1.97 |
|  | Cangzhou | 0.10 | 0.02 | 5.83 | 6.51 | 0.78 | 8.35 | 0.09 | 0.08 | 1.25 |
|  | Tianjin | 0.03 | 0.005 | 6.01 | 2.41 | 0.22 | 11.1 | 0.03 | 0.03 | 1.03 |
|  | Liaoyuan | 0.11 | 0.02 | 4.51 | 7.99 | 0.71 | 11.29 | 0.10 | 0.10 | 1.01 |
| YQ23 | Beijing | 0.03 | 0.002 | 20.17 | 2.58 | 0.26 | 9.97 | 0.03 | 0.03 | 0.85 |
|  | Urumchi | 0.08 | 0.02 | 4.72 | 5.96 | 0.61 | 9.91 | 0.08 | 0.06 | 1.54 |
|  | Cangzhou | 0.07 | 0.02 | 3.78 | 5.11 | 0.52 | 9.89 | 0.07 | 0.04 | 1.60 |
|  | Tianjin | 0.11 | 0.03 | 4.09 | 8.38 | 0.82 | 10.09 | 0.10 | 0.11 | 0.92 |
|  | Liaoyuan | 0.09 | 0.02 | 4.83 | 6.67 | 0.67 | 9.92 | 0.09 | 0.08 | 1.04 |
| DK301 | Beijing | 0.12 | 0.004 | 28.83 | 8.84 | 0.97 | 9.09 | 0.08 | 0.12 | 0.69 |
|  | Urumchi | 0.07 | 0.01 | 4.76 | 5.55 | 0.59 | 9.21 | 0.08 | 0.07 | 1.20 |
|  | Cangzhou | 0.07 | 0.02 | 4.01 | 4.78 | 0.55 | 8.67 | 0.06 | 0.06 | 1.15 |
|  | Tianjin | 0.11 | 0.02 | 6.06 | 7.84 | 0.58 | 13.91 | 0.10 | 0.10 | 1.03 |
|  | Liaoyuan | 0.07 | 0.02 | 3.99 | 5.2 | 0.65 | 8.04 | 0.07 | 0.06 | 1.20 |
| ZD958 | Beijing | 0.10 | 0.01 | 10.31 | 7.68 | 0.68 | 11.33 | 0.10 | 0.10 | 0.99 |
|  | Urumchi | 0.09 | 0.03 | 3.51 | 7.47 | 0.77 | 9.69 | 0.09 | 0.04 | 2.29 |
|  | Cangzhou | 0.13 | 0.03 | 5.05 | 9.36 | 0.84 | 12.2 | 0.12 | 0.13 | 0.97 |
|  | Tianjin | 0.10 | 0.02 | 4.15 | 6.91 | 0.82 | 8.30 | 0.09 | 0.08 | 1.20 |
|  | Liaoyuan | 0.10 | 0.02 | 5.65 | 7.17 | 0.47 | 15.35 | 0.09 | 0.09 | 0.98 |
| SEM |  | 0.006 | 0.002 | 1.131 | 0.431 | 0.043 | 0.366 | 0.005 | 0.006 | 0.083 |
| Regions |  |  |  |  |  |  |  |  |  |  |
|  | Beijing | 0.07<sup>b</sup> | 0.008<sup>c</sup> | 13.82<sup>a</sup> | 5.55<sup>b</sup> | 0.57 | 9.98 | 0.063<sup>b</sup> | 0.069<sup>bc</sup> | 1.02<sup>b</sup> |
|  | Urumchi | 0.09<sup>ab</sup> | 0.023<sup>a</sup> | 3.88<sup>b</sup> | 6.68<sup>ab</sup> | 0.70 | 9.66 | 0.093<sup>a</sup> | 0.055<sup>c</sup> | 1.88<sup>a</sup> |
|  | Cangzhou | 0.09<sup>ab</sup> | 0.019<sup>ab</sup> | 4.47<sup>b</sup> | 6.22<sup>ab</sup> | 0.64 | 9.99 | 0.083<sup>a</sup> | 0.074<sup>b</sup> | 1.19<sup>b</sup> |
|  | Tianjin | 0.08<sup>b</sup> | 0.017<sup>b</sup> | 4.92<sup>b</sup> | 6.15<sup>b</sup> | 0.57 | 11.37 | 0.078<sup>ab</sup> | 0.075<sup>b</sup> | 1.07<sup>b</sup> |
|  | Liaoyuan | 0.10<sup>a</sup> | 0.021<sup>a</sup> | 4.74<sup>b</sup> | 7.45<sup>a</sup> | 0.64 | 17.81 | 0.094<sup>a</sup> | 0.092<sup>a</sup> | 1.04<sup>b</sup> |
|  | SEM | 0.006 | 0.001 | 0.857 | 0.438 | 0.050 | 2.974 | 0.006 | 0.006 | 0.117 |
| P-value | Region | 0.048 | <.001 | <.001 | 0.047 | 0.28 | 0.02 | 0.005 | <.001 | <.001 |
|  | Cultivar | 0.01 | 0.01 | <.001 | 0.02 | 0.053 | 0.08 | 0.050 | 0.04 | 0.39 |
|  | Region × Cultivar | <.001 | <.001 | <.001 | <.001 | <.001 | <.001 | <.001 | <.001 | 0.26 |

Note: FKBN = Fengkenbainuo; YQ 889 = Yaqing 889; YQ 23 = Yuqing 23; DK 301 = Dongke 301; ZD958 = Zhengdan 958.

The results of CLA and PCA analyses were presented in Figure 1. It was evident from Figure 1 that the protein structural makeup of FKBN and DK301 cultivated in Tianjin were distinguished from those grown in other regions. As for cultivars of YQ889 and YQ23, clear separate ellipses were found between Cangzhou and other growing locations. For the remaining cultivars, heavy overlaps were observed for the protein primary structural data. These results indicated that for some WPSC cultivars, internal protein molecular makeups were altered when environmental conditions changed, which was partly consistent with our data from univariate spectral analysis (Table 5).

**Fig. 1.**
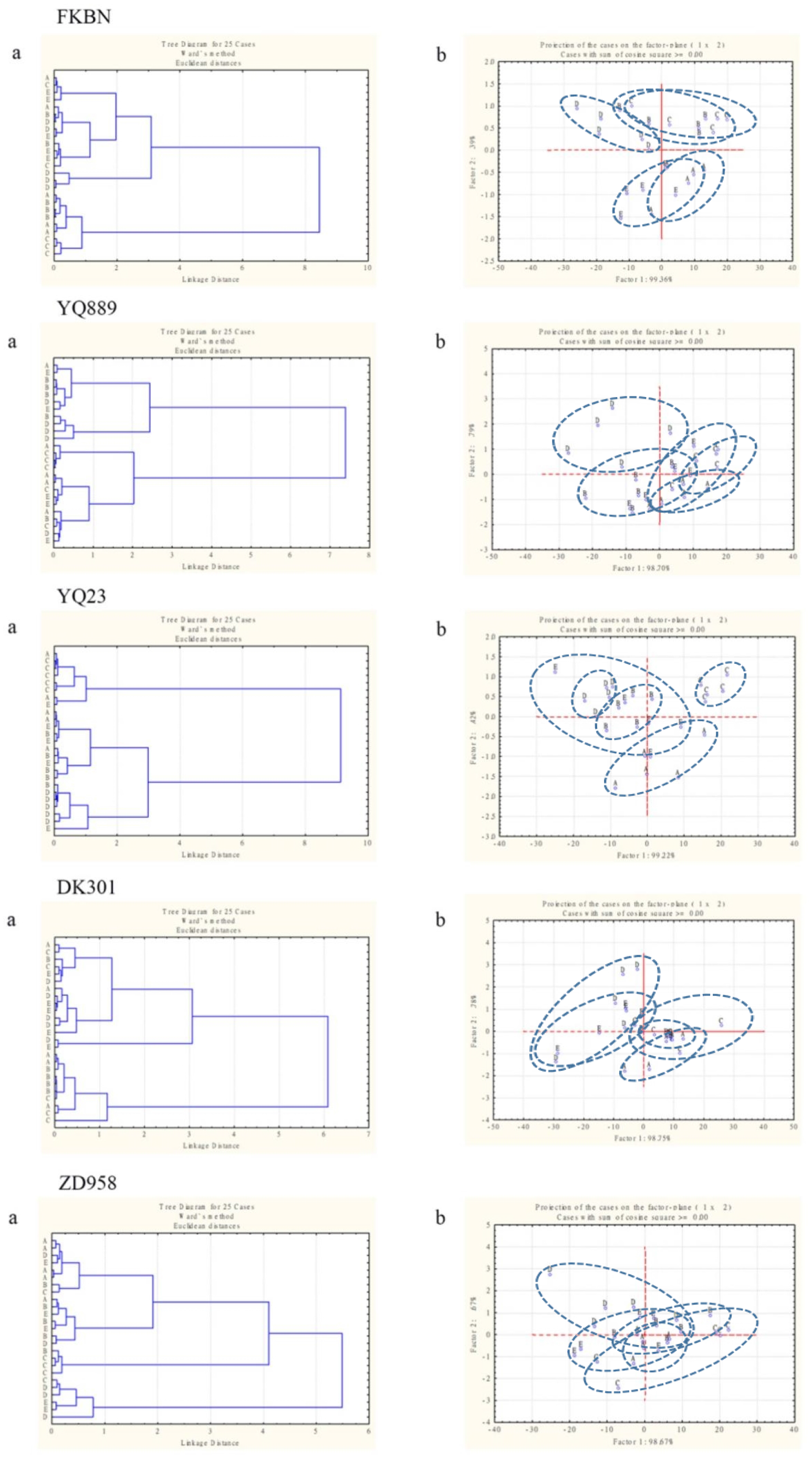
Multivariate molecular spectral analyses of protein molecular spectral feature in protein amide I and amide II fingerprint region (ca. 1712−1489 cm^−1^) of various whole-plant silage corn cultivars from different regions. a: CLA analysis; b: PCA analysis; A = Beijing; B = Urumchi; C = Cangzhou; D = Tianjin; E = Liaoyuan

### Correlation of protein molecular structural features with protein chemical profiles as well as in situ degradation parameters of whole plant silage corn

Figure 2 visualized results of the relationship between protein molecular structural features and CP chemical profiles, CNCPS subfractions and *in situ* rumen degradation characteristics. The area ratio of amide I to II was negatively correlated with the contents of SCP (*δ* = -0.66; *P* = 0.002), CP (*δ* = -0.61; *P* = 0.006), NPN (*δ* = -0.56; *P* = 0.004), ADICP (*δ* = -0.43; *P* = 0.008) and PA_1_(*δ* = -0.38; *P* = 0.047), whereas positively correlated with PB_1_ (*δ* = 0.58; *P* = 0.01). The *α*-helix to *β*-sheet height ratio was positively correlated with PB_2_ (*δ* = 0.22; *P* = 0.048) while negatively correlated with PB_1_ (*δ* = -0.46; *P* = 0.002). Similarly, the correlation between protein molecular structures and chemical profiles was demonstrated by previous study [44]. However, Liu et al. [45] found negative relationships between *α*-helix to *β*-sheet height ratio with PB_2_ (*δ* = -0.45; *P* < 0.01) in wheat, corn and triticale grains, which were not in line with current study.

**Fig. 2.**
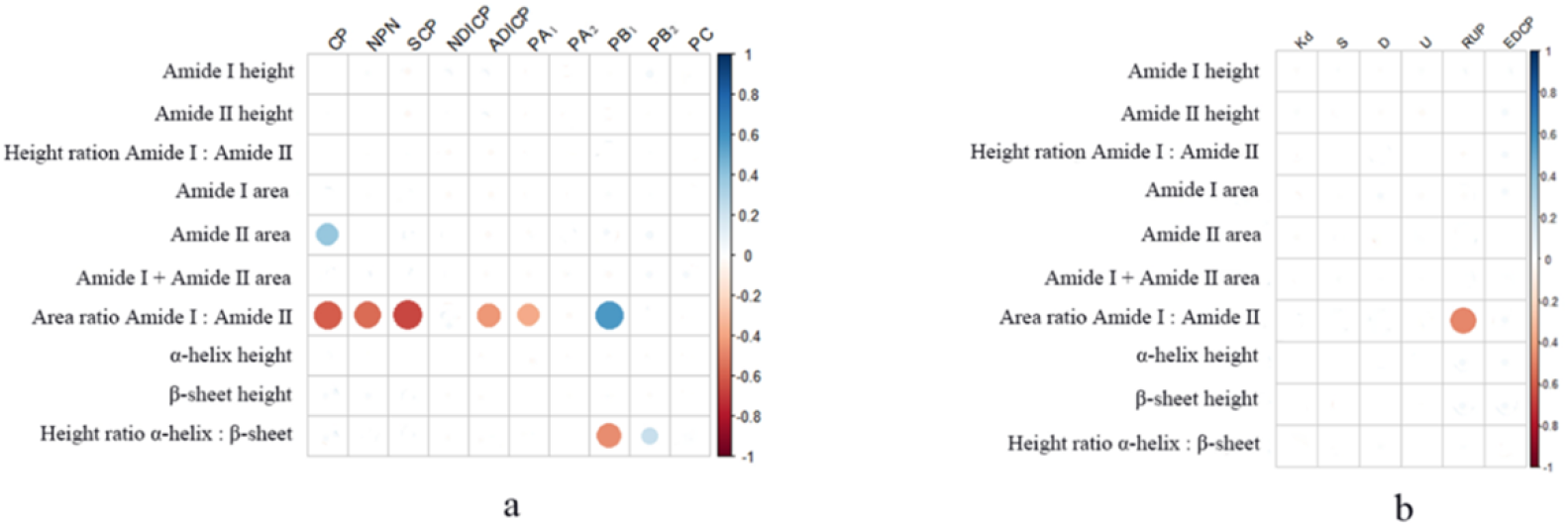
Correlation of protein molecular structural features with chemical profile and CNCPS subfractions and in situ biodegradation parameters. a, correlation of protein molecular structure features with protein chemical profile and CNCPS subfractions; b, correlation of protein molecular structure features with in situ biodegradation parameters. CP, crude protein; NPN, non-protein nitrogen; SCP, soluble CP; NDICP, neutral detergent insoluble CP; ADICP, acid detergent insoluble CP; PA_1_, ammonia; PA_2_, soluble non-ammonia CP; PB_1_, moderately degradable CP; PB_2_, slowly degradable CP; PC, unavailable CP; K_d_, ruminal degradation rate; S, soluble CP in the in situ incubation; D, insoluble but potentially degradable CP in the in situ incubation; U, potential undegradable CP in the in situ incubation; RUP, the CP fraction of bypass rumen; EDCP, the effective degradable fraction of CP in the rumen.

The amide I to II area ratio had a negative correlation with RUP content (*δ* = -0.48; *P* = 0.02). The relations of spectral characteristics with *in situ* CP ruminal degradation kinetics were not significant. Numerous studies have reported correlations between protein molecular structural traits and ruminal degradation parameters in varieties of feedstuff [46-47]. For instance, Zhang and Yu [46] represented relationships of amide I area (*δ* =0.97; *P* = 0.005), amide II area (*δ* = 0.90; *P* = 0.036), *α*-helix height (*δ* = 0.96; *P* = 0.011), *β*-sheet height (*δ* = 0.98; *P* = 0.004), in conjunction with *α*-helix to *β*-sheet height ratio (*δ* = -0.99; *P* = 0.00) and RDP in hulless barley and bioethanol coproducts of wheat or dried distillers grains with solubles. However, Gomaa et al. [36] suggested that no relationship was observed between area ratio of amide I to II and RUP in canola seeds. Although the data was very limited, these inconsistent results showed that the correlations between protein molecular structural characteristics and CP chemical profiles as well as rumen degradable characteristics were specific to some extent.

## Conclusion

In conclusion, there were significant interaction between growing region and WPSC cultivar for most of the measured morphological characteristics, CP chemical profiles, CNCPS subfractions, *in situ* ruminal degradation parameters, and protein molecular structural features. In detail, our study showed that even for the same cultivar, the protein nutritional values and molecular structures significantly varied between growing regions. Besides, the between-region variations in protein chemical profiles, CNCPS subfractions and *in situ* degradation parameters were strongly correlated with some protein molecular spectral parameters. Further in-depth studies are required to establish some predictive models based on novel technology combined with reliable algorithm for estimating nutritional contents and rumen degradable characteristics.

## Acknowledgments

The authors would like to thank supplementation from Northeast Agricultural University.

